# C-di-GMP signalling links biofilm formation and Mn(II) oxidation in *Pseudomonas resinovorans*

**DOI:** 10.1101/2022.07.20.500916

**Authors:** Ainelen Piazza, Lucila Ciancio Casalini, Federico Sisti, Julieta Fernández, Jacob G. Malone, Jorgelina Ottado, Diego O. Serra, Natalia Gottig

## Abstract

Bioaugmentation of biological sand filters with Mn(II)-oxidizing bacteria (MOB) is used to increase Mn removal efficiencies from groundwater. While the biofilm-forming ability of MOB is important to achieve optimal Mn filtration, the regulatory link between biofilm formation and Mn(II) oxidation remains unclear. Here, the environmental isolate *P. resinovorans* strain MOB-513 was used as a model to investigate the role of c-di-GMP, a second messenger crucially involved in the regulation of biofilm formation by *Pseudomonas*, in the oxidation of Mn(II). A novel role for c-di-GMP in the up-regulation of Mn(II) oxidation through induction of the expression of Manganese-Oxidizing Peroxidase (MOP) enzymes was revealed. MOB-513 macrocolony biofilms showed a strikingly stratified pattern of Mn oxides (BMnOx) accumulation in a localized top layer. Remarkably, elevated cellular levels of c-di-GMP correlated not only with increased accumulation of BMnOx in the same top layer, but also with the appearance of a second BMnOx stratum in the bottom region of macrocolony biofilms and the expression of *mop* genes correlated with this pattern. Proteomic analysis under Mn(II) conditions revealed the up-regulation of a GGDEF/EAL-domain protein and a PilZ-domain protein, providing a molecular link between c-di-GMP signalling and Mn(II) oxidation. Finally, we considered the biotechnological relevance of understanding the role of c-di-GMP in MOB-513 and observed that high c-di-GMP levels are correlated with higher lyophilisation efficiencies and higher groundwater Mn(II) oxidation capacity of lyophiles. Advancing understanding of these mechanisms is essential to improve the biotechnological application of bacterial inocula designed for removing Mn in biological filter systems.

**IMPORTANCE:** The presence of Mn(II) in groundwater - a common source of drinking water-is a cause of water quality impairment, interfering with its disinfection, causing operation problems and affecting human health. Purification of groundwater containing Mn(II) plays an important role in environmental and social safety. The typical method for Mn(II) removal is based on bacterial oxidation of metals to form insoluble oxides that can be filtered out of the water. Evidence of reducing the start-up periods and enhancing Mn removal efficiencies through bioaugmentation with appropriate biofilm-forming and MOB has emerged. As preliminary data suggest a link between these two phenotypes in *Pseudomonas* strains, the need to investigate the underlying regulatory mechanisms is apparent. The significance of our research lies in determining the role of c-di-GMP for increased biofilm-formation and Mn(II)-oxidizing capabilities in MOBs, which will allow the generation of super biofilm-elaborating and Mn-oxidizing strains, enabling their implementation in biotechnological applications.

## INTRODUCTION

The presence of soluble manganese Mn(II) affects the quality of groundwater, a source of drinking water for many populations, and is an important environmental concern (1–3). Biological sand filter technology, based on bacterial oxidation of metals to form insoluble oxides that can be filtered out of the water, is widely used for groundwater potabilization. Bioaugmentation of this process through the inoculation of sand filters with appropriate Mn(II)-oxidizing bacteria (MOB) optimizes Mn removal (4–8).

Biofilms are sessile and densely populated communities of bacterial cells, surrounded by an extracellular matrix (ECM) that typically contains exopolysaccharides, proteins or extracellular DNA and serves as a shield that protects the cells from external aggressions and stresses (9). The biofilm-forming capability of bacteria is important to achieve the optimal filtration of groundwater for metals, as they have to be oxidized and retained into the biofilter matrix surface, and to decrease the loss of the MOBs, which can otherwise be washed out of the system (4, 5, 8). It has been recently shown that powdered MOB inoculates prepared by vacuum lyophilisation are useful to inoculate sand filters and remove Mn with high efficiencies (10). Moreover, lyophilisation efficiencies increase when MOB are grown under biofilms instead of shaking culture conditions (10), demonstrating that an understanding of biofilm formation in MOBs may help to implement these bacteria in biotechnological applications.

Previous work with the environmental isolate *Pseudomonas resinovorans* MOB-513 has shown that the Mn(II) oxidation phenotype in this bacterium is biofilm dependent (11). Accordingly, in *P. putida* GB-1, Mn(II) oxidation was associated with the sessile lifestyle, and influenced by flagella synthesis and by the contact with surface (12). These results suggest a correlation between Mn-oxidizing phenotype and biofilm growth.

In *Pseudomonas spp*., the main regulator of biofilm formation is the second messenger bis-(3′-5′)-cyclic dimeric guanosine monophosphate (c-di-GMP). This ubiquitous signalling molecule controls the transition of bacteria from a motile to a sessile lifestyle and vice versa. In almost all cases, high cellular levels of c-di-GMP promote biofilm formation, while low c-di-GMP levels stimulate bacterial motility and often biofilm dispersal (13). The cellular pool of c-di-GMP is regulated by GGDEF-domain-containing diguanylate cyclases (DGCs) that synthesize the messenger and EAL/HD-GYP-domain-containing phosphodiesterases (PDEs) that degrade it (13). c-di-GMP binds to an array of intracellular receptors (RNA riboswitches, PilZ domains, degenerate GGDEF/EAL domains and numerous other protein folds) that go on to exert specific cellular effects at transcriptional, translational and post-translational levels (14, 15).

The bacterial enzymes involved in Mn(II) oxidation (Mn oxidases) that have been characterized so far belong to two families of proteins: the multicopper oxidases (MCOs) and manganese peroxidases (MOPs) (16). MCO enzymes oxidize organic compounds using copper as a cofactor and have a role in metal homeostasis and in different biosynthetic pathways (17). *P. putida* GB-1 has two MCO-type enzymes with Mn oxidase activities, MnxG and McoA (18). The expression of *mnxG* and *mcoA* genes is positively regulated by a two-component regulatory pathway, although the external signals inducing the system remain unclear (19). The MOP enzyme MopA, implicated in directly oxidizing Mn(II), was also found in *P. putida* GB-1 (20). In this bacterium, the deletion of the transcriptional regulator *fleQ* results in the over-production and secretion of MopA, while the activity of both MnxG and McoA decrease (20). FleQ is a c-di-GMP-responsive transcription factor, which binds c-di-GMP as a function of the cellular levels of this messenger, to regulate flagella synthesis and biofilm formation (21). This suggests not only that different Mn(II) oxidases function in planktonic and biofilm cells (20), but also implies a connection between c-di-GMP signalling and the Mn(II) oxidation process.

Collectively, this led us to hypothesize that c-di-GMP and its associated signalling network is the regulatory link between Mn(II) oxidation and biofilm formation in *Pseudomonas*. Therefore, MOB-513 was used as a bacterial model to investigate for the first time how variations in c-di-GMP levels-achieved through the ectopic expression of DGCs or PDEs-influence Mn(II) oxidation. Specifically, we determined how c-di-GMP contributes to Mn(II) oxidation in MOB-513, demonstrating a novel role for this messenger in the up-regulation of this process and highlighting the feasibility of generating hyper-oxidizer bacterial strains to optimise Mn removal from groundwater.

## RESULTS

### Analysis of the MOB-513 genome reveals several genes potentially involved in Mn(II) oxidation and c-di-GMP metabolism

The complete MOB-513 genome sequence was analysed (Fig. S1) to gain a better insight into the mechanism of Mn(II) oxidation and c-di-GMP metabolism. We identified potential Mn(II) oxidases by comparison to experimentally verified MnxG, McoA and MopA enzymes from *P. putida* GB-1 (Table 1). This analysis revealed two genes, *Pres513_3924* and *Pres513_3296*, homologous to *mnxG* and *mcoA*, respectively. Moreover, three genes homologous to *mopA* were detected: *Pres513_7013, Pres513_7014* and *Pres513_5806*. (Table 1). Additionally, we identified 21 proteins containing the GGDEF domain, *i*.*e*., putative DGCs, and 14 proteins containing both GGDEF and EAL domains.

**Table 1.** List of genes in strain MOB-513 with sequence homology to putative Mn(II) oxidases from *P. putida* GB-1.

| Gene | Locus tag MOB-513 | Predicted function | % Identity | % Similarity | E value |
| --- | --- | --- | --- | --- | --- |
| *mnxG* | *Pres513_3924 (mco3924)* | hypothetical protein | 74% | 83% | 0.0 |
| *mcoA* | *Pres513_3296 (mco3296)* | hypothetical protein | 31% | 45% | 2e<sup>-76</sup> |
| *mopA* | *Pres513_7013*<br>*(mop7013)* | RTX toxins and<br>related Ca<sup>2+</sup>-binding<br>proteins | 55% | 66% | 2e<sup>-11</sup> |
| *mopA* | *Pres513_7014*<br>*(mop7014)* | RTX toxins and<br>related Ca<sup>2+</sup>-binding<br>proteins | 52% | 63% | 3e<sup>-20</sup> |
| *mopA* | *Pres513_5806*<br>*(mop5806)* | RTX toxins and<br>related Ca<sup>2+</sup>-binding<br>proteins | 65% | 74% | 3e<sup>-13</sup> |

### Overexpression of *BdcB* or *BpdeA* in MOB-513 changes intracellular c-di-GMP levels, inversely regulating biofilm formation and swimming motility

The large number of genes encoding putative DGCs and/or PDEs found in MOB-513 suggests a complex c-di-GMP signalling network operates in this strain, making it difficult to predict which of these enzymes act to globally regulate the cellular pool of c-di-GMP. Thus, to increase and decrease the intracellular levels of c-di-GMP in MOB-513, we opted to ectopically overexpress *BdcB* and *BpdeA*, well-characterized DGC and PDE genes from *B. bronchiseptica*, respectively (22, 23). MOB-513 overexpressing *BdcB* (MOB-513-pBdcB) presented significantly higher c-di-GMP levels than the MOB-513 strain harbouring the pEmpty vector (MOB-513-pEmpty); while inversely, MOB-513 overexpressing *BpdeA* (MOB-513-pBpdeA) showed significantly lower c-di-GMP levels than MOB-513-pEmpty (Fig. 1A). These results are additionally supported by measurements of [c-di-GMP] in *P. aeruginosa* PAO1 overexpressing the DGC gene *wspR*, previously shown to overproduce c-di-GMP and in the corresponding control strain (24) (Fig. 1A).

**FIG 1.**
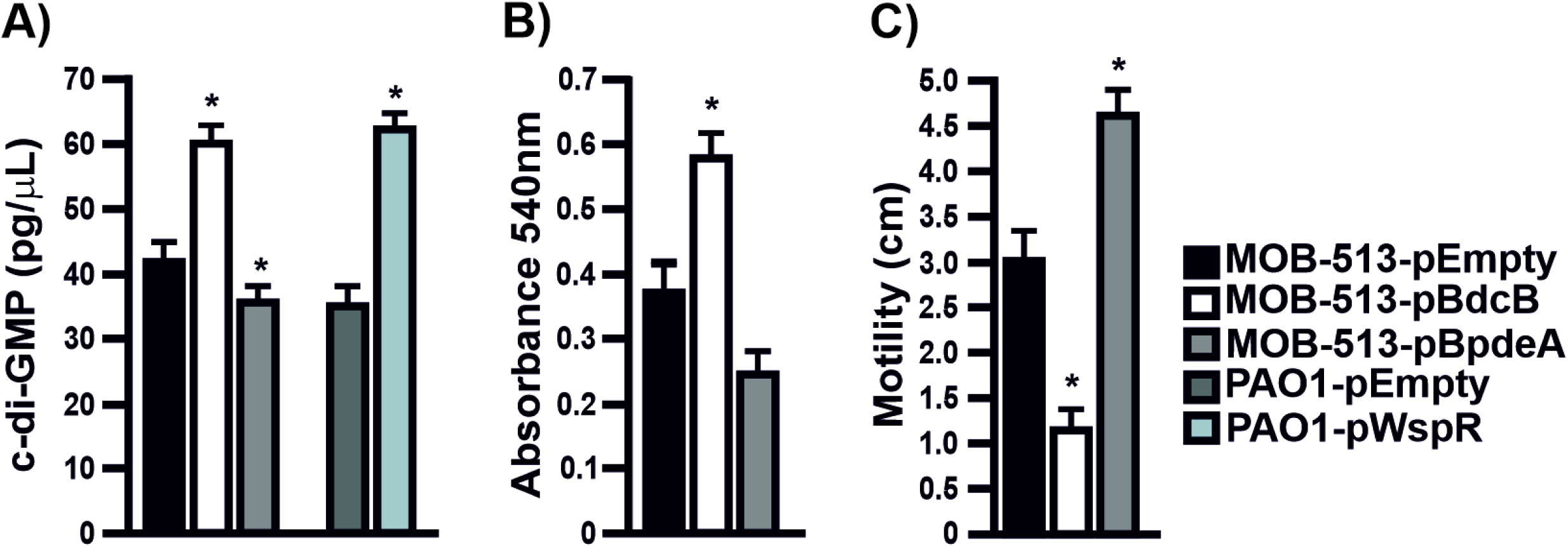
c-di-GMP levels, biofilm formation and swimming motility in MOB-513 strains overexpressing *BdcB* or *BpdeA*. (A) c-di-GMP concentration in cells of *P. resinovorans* MOB-513 and *P. aeruginosa* PAO1 strains. c-di-GMP levels were quantified using the Cyclic-di-GMP Assay Kit (Lucerna). *P. aeruginosa* strains PAO1-pWspR and PAO1-pEmpty were used as controls for the assays. (B) Biofilm formation by MOB-513 strains at the air-liquid interface assessed by the crystal violet (CV) assay. (C) Migration zone diameters of the MOB-513 strains grown on LB swimming plates for 72 h at 28°C. Quantifications were performed in triplicate and mean values ± SD are presented. Data were statistically analysed using one-way ANOVA and asterisks indicate significant differences compared to the controls (p<0.05).

To corroborate the physiological effects of raising and lowering c-di-GMP levels in MOB-513, biofilm formation and motility-two bacterial phenotypes shown to be inversely regulated by c-di-GMP in other bacteria (13)-were analysed. As expected, MOB-513-pBdcB showed an about 1.5-fold increase in biofilm biomass accumulation and a 2.8-fold decrease in swimming motility relative to MOB-513-pEmpty (Fig. 1B-C). Conversely, in MOB-513-pBpdeA a 1.4-fold reduction in biofilm biomass and about a 1.5-fold increase in swimming motility compared to the control strain was observed (Fig. 1B-C).

### Changes in c-di-GMP levels strongly influence the onset and performance of Mn(II) oxidation by MOB-513 in biofilms

In large colony biofilms-known as macrocolonies (25)-the formation of BMnOx by MOBs can be readily visualized by the appearance of brown colour when these biofilms are grown on Lept agar medium supplemented with Mn(II) (11). Thus, we used this biofilm model system to evaluate the effect of increasing or decreasing the cellular levels of c-di-GMP on the Mn(II) oxidation performance of MOB-513. As shown in Fig. 2A and consistent with previous observation (11), in the presence of Mn(II), macrocolonies of the MOB-513 control strain started to develop a light brown coloration by day 3, which became much more intense and visible by day 4 and 5 when the biofilms reached their maximum expansion. This phenotype was strictly dependent on the presence of Mn(II) in the medium. Remarkably, in macrocolonies of MOB-513-pBdcB the brown colour appeared already by day 2 (Fig. 2), indicating that increased c-di-GMP levels accelerate the onset of BMnOx formation. Conversely, in macrocolonies of MOB-513-pBpdeA the appearance of BMnOx was delayed, starting at day 4 and becoming readily visible by day 5 (Fig. 2). This opposite effect reinforces the involvement of c-di-GMP in regulating the onset of Mn(II) oxidation by MOB-513.

**FIG 2.**
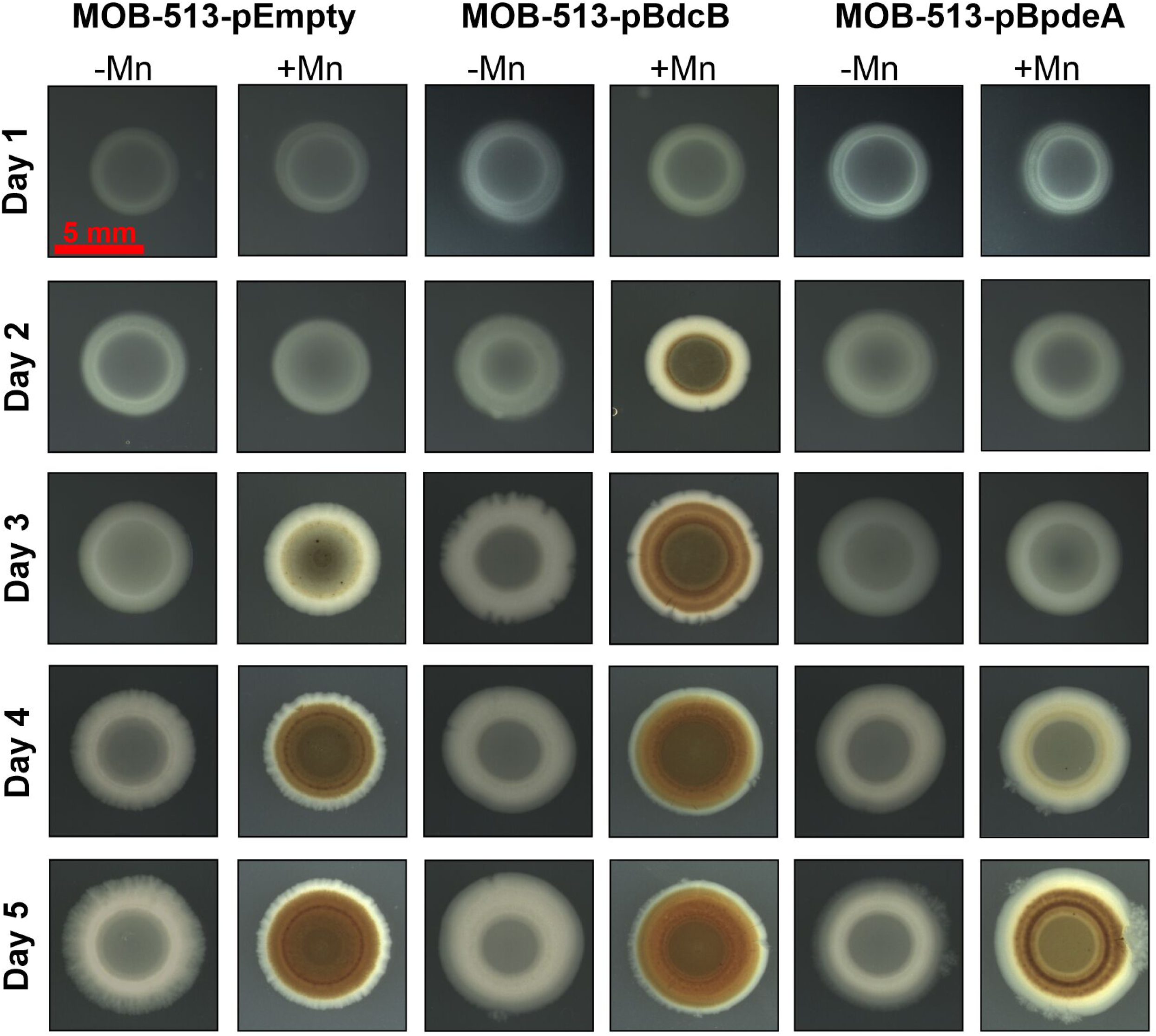
Time course of Mn(II) oxidation in macrocolony biofilms of MOB-513 strains. Top view of MOB-513-pEmpty, MOB-513-pBdcB, and MOB-513-pBpdeA macrocolonies at distinct stages of growth, exhibiting (or not) Mn(II) oxidation phenotypes (brown colour). Macrocolonies were set on Lept (-Mn) and Lept-Mn (+Mn) plates, incubated at 28°C and imaged daily for 5 days. The red scale bar represents 5 mm.

Quantitative analysis of BMnOx accumulation and Mn-oxidase activity assays over time in macrocolony biofilms of the three MOB-513 strains were performed using the LBB assay. Elevated cellular levels of c-di-GMP correlated with early and increased BMnOx production in macrocolonies (about 1.6-fold the amount of BMnOx produced by the control MOB-513-pEmpty strain at day 5), while decreased cellular levels of c-di-GMP led to delayed and diminished BMnOx production (about half of the BMnOx yielded by the control MOB-513-pEmpty strain at day 5). These effects are related specifically to changes in the cellular levels of c-di-GMP as macrocolonies of the three MOB-513 strains showed essentially the same growth pattern as determined by viable cell counting (Fig. 3A).

**FIG 3.**
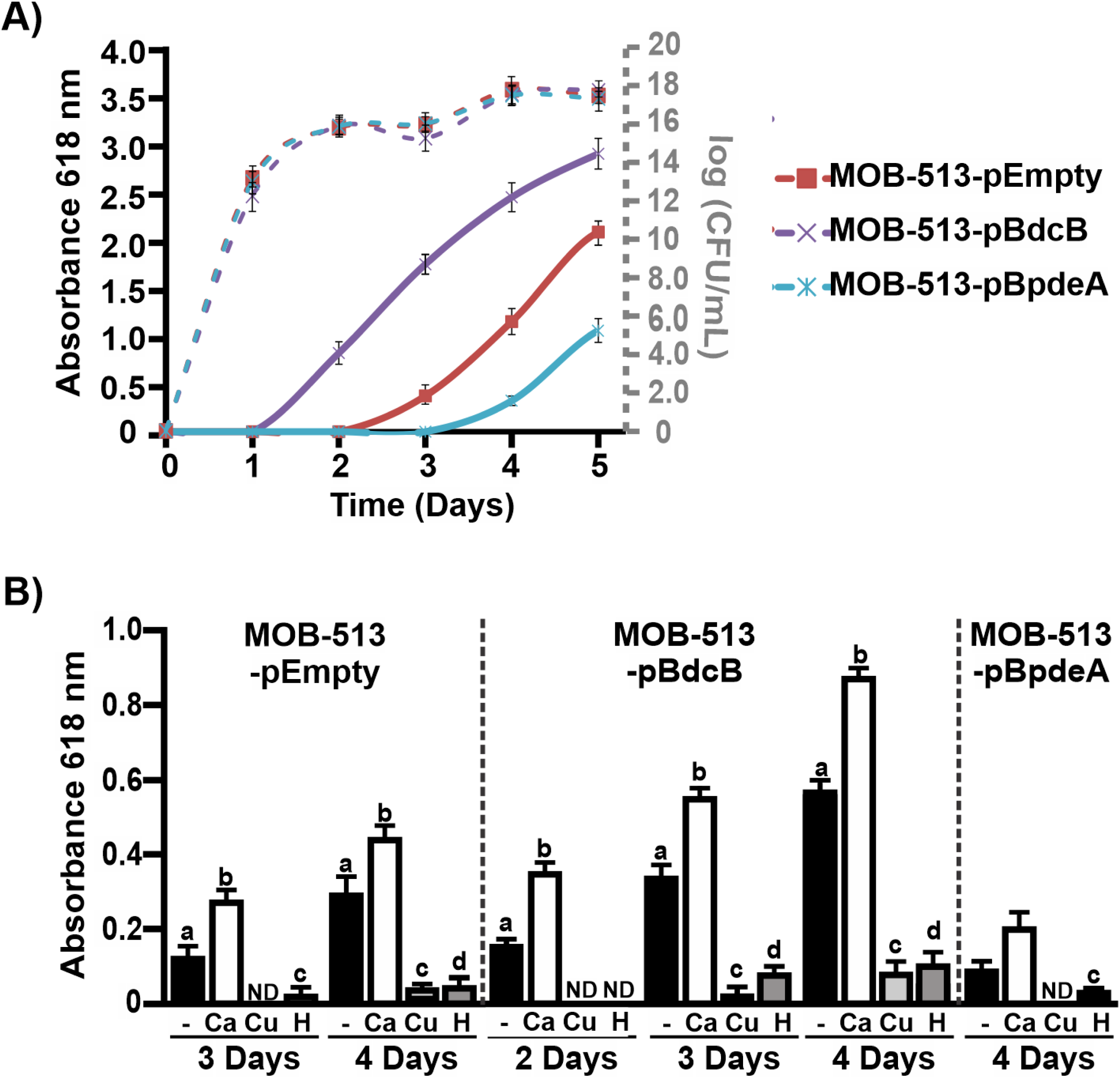
Involvement of c-di-GMP signalling in Mn(II) oxidation by MOB-513 strains. (A) Mn(II) oxidation capacities of MOB-513-pEmpty, MOB-513-pBdcB and MOB-513-pBpdeA strains. Quantification of BMnOx formed by the strains was assessed daily for 5 days using the LBB assay. Axis on the left side presents values of absorbance at 618 nm as determination of BMnOx (solid lines). Axis on the right side of the plot (grey colour) presents log CFU/mL values as determination of growth (dashed lines). Absorbance and log CFU/ml values represent the mean of three biological replicates and three technical replicates at each time point analysed. Error bars indicate the SD. Data were statistically analysed using one-way ANOVA (p<0.05). (B) *In vitro* Mn(II) oxidase activities. Mn(II) oxidase activities were assayed in total protein extracts obtained from MOB-513-pEmpty, MOB-513-pBdcB, and MOB-513-BpdeA macrocolonies grown on Lept-Mn for 2, 3 and 4 days. Quantifications were performed from three biological replicates and three technical replicates per sample. Mean values and SD are presented. Data were statistically analysed using one-way ANOVA followed by Tukey’s test. Bars with different letters a, b, c, and d indicate significant differences between treatments (p<0.05). ND indicates that no activity was detected.

Mn-oxidase activities correlated with BMnOx production and no evidence of Mn-oxidase activity in heat-treated control samples (p<0.05) was detected, demonstrating that Mn(II) oxidation occurs in MOB-513 through enzymatic processes (Fig. 3B). Moreover, two metal ions were tested for their effects on enzymatic activity, Ca(II) and Cu(II), which enhance MopA (26) and MCO activities (27), respectively. Only the presence of Ca(II) significantly enhanced the Mn(II) oxidizing activities (Fig. 3B), suggesting the involvement of MOPs in the Mn(II) oxidation process in MOB-513.

Altogether, these results not only confirmed that changes in the cellular levels of c-di-GMP altered the onset of Mn(II) oxidation, but also showed that such changes strongly influenced the overall yield of BMnOx produced by inducing shifts in enzymatic activity.

### BMnOx accumulation occurs in well-defined and restricted zones of MOB-513 biofilms in a c-di-GMP-dependent manner

While in macrocolony biofilms brown coloration visually denotes the presence of BMnOx, it does not precisely reveal how the oxidized mineral distributes across the internal section of the biofilm. To gain more insights into this aspect, macrocolony biofilms of MOB-513 strains grown on Lept or Lept-Mn were cross-sectioned and microscopically examined for the presence of BMnOx. In MOB-513-pEmpty, BMnOx started to be detected by day 3 in a localized manner in the upper part of the biofilm (Fig. 4A). Interestingly, by day 4 the amount of BMnOx increased and was strikingly accumulated as a well-defined dark band near the macrocolony surface (Fig. 4B). The same spatial pattern of BMnOx distribution was observed in cross-sections of macrocolonies of MOB-513-pBdcB, but at day 3, *i*.*e*., one day earlier (Fig. 4A). Remarkably, at day 4, MOB-513-pBdcB macrocolonies exhibited further accumulation of BMnOx in a second layer at the bottom of the biofilm, close to the nutrient-providing agar and remote from the colony interface with the air (Fig. 4B). In contrast, at day 4 macrocolonies of MOB-513-pBpdeA showed very little BMnOx in the upper part of the biofilm section (Fig. 4B). The second band of BMnOx found in MOB-513-pBdcB macrocolonies was not observed in MOB-513-pEmpty or MOB-513-pBdpeA macrocolonies at any time point assayed. This explains the larger amount of BMnOx detected in MOB-513-pBdcB macrocolonies in the LBB assay compared with macrocolonies of the two other strains. As expected, no accumulation of BMnOx was observed when macrocolonies of the three strains were grown in the absence of Mn(II) (Fig. 4C).

**FIG 4.**
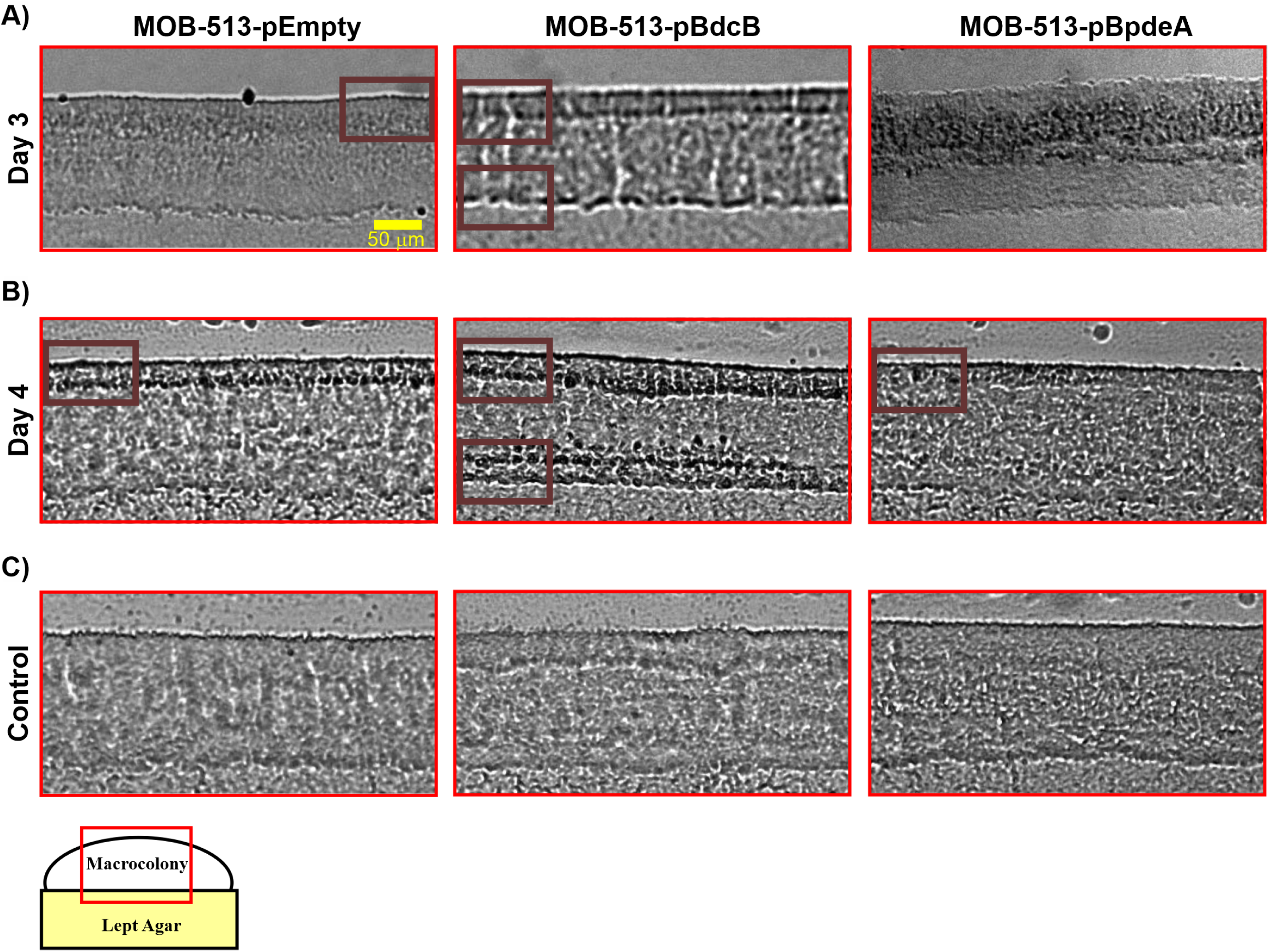
Cryosectioning and brightfield microscopy approach to analyse BMnOx depositions. (A) and (B) Images of 5-μm-thin vertical sections of 3- and 4-day-old macrocolonies of MOB-513-pEmpty, MOB-513-pBdcB and MOB-513-pBpdeA strains, respectively grown on Lept-Mn medium. Dark areas (Bordeaux boxed) in the images correspond to BMnOx deposited in the top layer of the macrocolonies for all the strains and a second layer of BMnOx located at the bottom of the macrocolony in MOB-513-pBdcB. (C) Images of 5-μm-thin vertical sections of 4-day-old macrocolonies of all strains grown in the absence of Mn(II) in the Lept medium. Representative images of phenotypes at day 4 are shown here, but the same results were observed for all the times assayed.

### The expression of MOPs is induced by Mn(II) and c-di-GMP in MOB-513 macrocolonies

To understand which Mn(II) oxidases are activated in the different MOB-513 macrocolony biofilms, the transcriptional profiles of *mco3924, mco3296, mop7013, mop7014* and *mop5806* genes (Table 1) were analysed. The expression of *mco3924, mco3296* and *mop5806* were unaffected under biofilm conditions in any strain in the presence of Mn(II) at any of the assayed times (Fig. S2 and Fig. 4A-C). However, *mop7013* and *mop7014* were differentially expressed. In MOB-513-pEmpty, both genes showed increased expression in the presence of Mn(II) from day 3 (p<0.05) (Fig. 5A). In MOB-513-pBdcB, *mop7013* and *mop7014* mRNA abundance was induced earlier (day 2) and increased relative to MOB-513-pEmpty (p<0.05) (Fig. 5B). In MOB-513-pBpdeA, a reduced and delayed induction of both genes compared with MOB-513-pEmpty was observed (p<0.05) (Fig. 4C). The timing of *mop7013* and *mop7014* gene expression correlates with the appearance of BMnOx and Mn-oxidase activity in the tested strains (Fig. 2), suggesting that Mop7013 and Mop7014 are the main enzymes involved in Mn(II) oxidation in MOB-513 macrocolony biofilms. These two predicted peroxidases share a conserved domain architecture with the characterized MopA protein from *A. manganoxydans* strain SI85-9A1 and *P. putida* GB-1 (20, 26) (Fig. S3).

**FIG 5.**
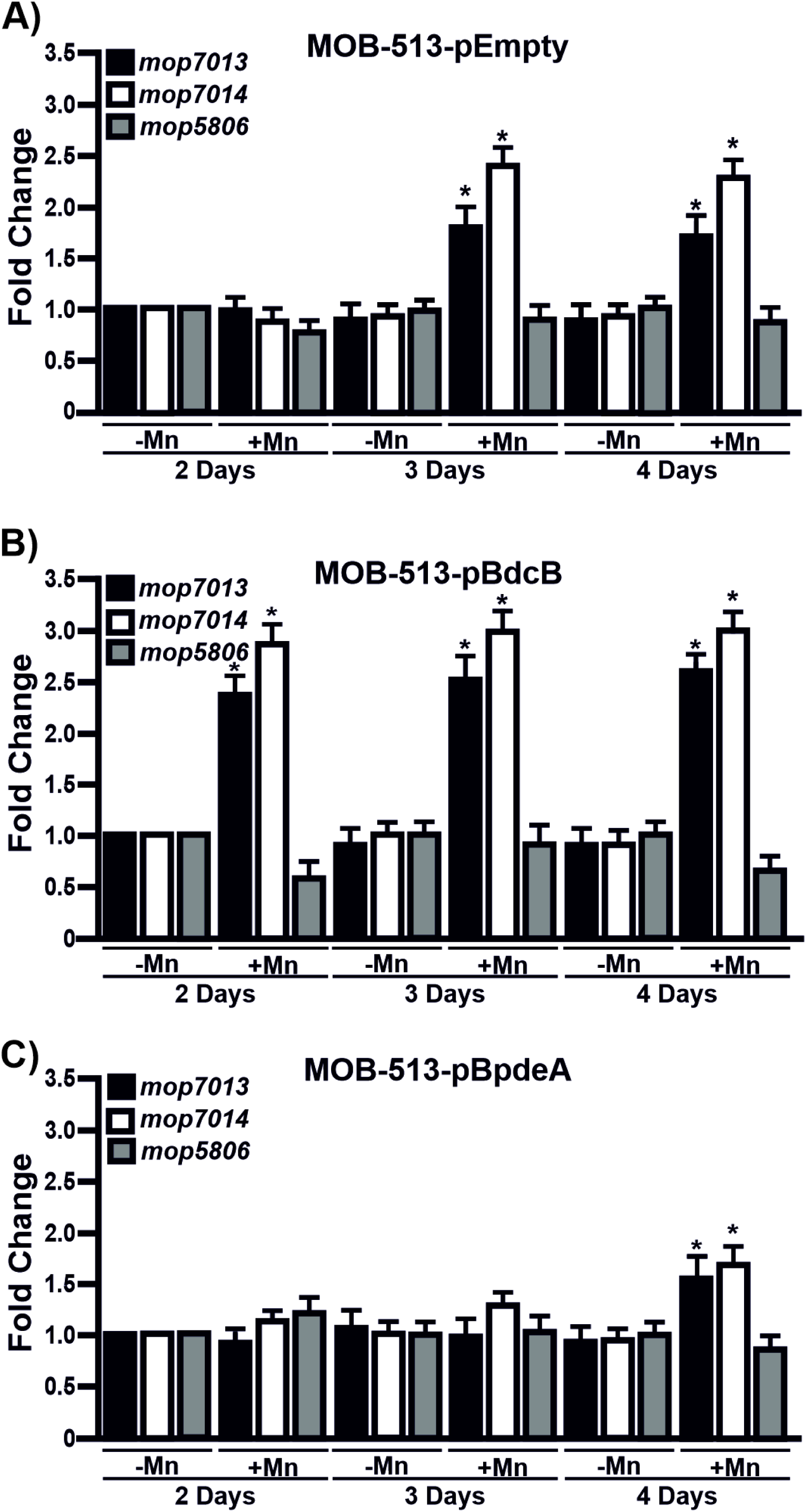
Expression of *mop* genes in MOB-513 strains. The macrocolonies of the strains MOB-513-pEmpty (A), MOB-513-pBdcB (B) and MOB-513-pBpdeA (C) were grown on Lept and Lept-Mn and subjected to qRT-PCR assays to analyse the expression of *mop7013, mop7014* and *mop5806*. The *rpoD* gene was used as internal control for the calculation of relative gene expression. Bars indicate the expression levels of the genes in Lept-Mn relative to the expression levels in the absence of Mn(II). Values are the means of three biological replicates with three technical replicates each. Error bars indicate standard deviation. Data were analysed by Student’s t-test and asterisks indicate significant differences (p<0.05) between samples grown in the presence and absence of Mn(II).

### *mop* gene expression spatially correlates with BMnOx production in MOB-513-pBdcB macrocolonies

Macrocolony biofilms represent a highly structured type of biofilm with a clear stratification of metabolic activities (28, 29). In the MOB-513-pBdcB macrocolony biofilm, the upper BMnOx layer is further away from the nutrient-providing agar and exposed to a higher oxygen concentration than the bottom BMnOx layer, which is right above the agar surface. Therefore, Mn(II) oxidation may be mediated by different enzymatic activities in each BMnOx-producing cell layers. To study if *mop7013* and *mop7014* are actively expressed in these specific cell layers, *gfp* reporter fusions to their predicted promoters (Fig. S3A) were constructed to analyse their spatial expression patterns in MOB-513-pBdcB macrocolony biofilms. As a control, a *gfp* reporter fusion to the reference gene *rpoD* promoter was also constructed. Macrocolonies of MOB-513-pBdcB transformed with these reporter fusions and the empty pPROBE-KT vector (control) (Table 3), were grown on Lept and Lept-Mn. Gfp fluorescence was detected only in MOB-513-pBdcB-pMop7013 and MOB-513-pBdcB-pRpoD, while no fluorescence was detected for MOB-513-pBdcB-pMop7014 or MOB-513-pBdcB-pPROBE-KT (Fig. S4). In addition, *gfp* expression is induced in the presence of Mn(II) in MOB-513-pBdcB-pMop7013 but not in MOB-513-pBdcB-pRpoD (Fig. S4). Taken together, these results suggest that *mop7013* and *mop7014* expression depends on the promoter found upstream of *mop7013* and that the putative promoter predicted for *mop7014* is not functional (possibly for technical reasons) or has a role as a regulatory element of *mop7014*.

Next, 3-day-old macrocolonies of the strains were thin-sectioned and the resulting sections examined with fluorescence microscopy (Fig. 6). In agreement with our Gfp fluorescence quantifications (Fig. S4), cryosections of the MOB-513-pBdcB-pMop7014 and MOB-513-pBdcB-pPROBE-KT strains did not show fluorescence signals (data not shown). In the absence of Mn(II), MOB-513-pBdcB-pMop7013 showed a homogenous distribution of Gfp expression across the entire section of the macrocolony (Fig. 6 and Fig. S5). In the presence of Mn(II), an increased fluorescence signal was observed (Fig. 6 and Fig. S5) but the fluorescence was not uniform throughout the macrocolony biofilm. A higher Gfp expression in the upper and bottom cell layers was observed, correlating with the spatial pattern of BMnOx produced by the strain. In MOB-513-pBdcB-pRpoD macrocolony cross-sections, the intensity and spatial distribution of the Gfp fluorescence signal was similar in the presence or absence of Mn(II), being relatively homogenous throughout the macrocolony section (Fig. 6 and Fig. S5).

**FIG 6.**
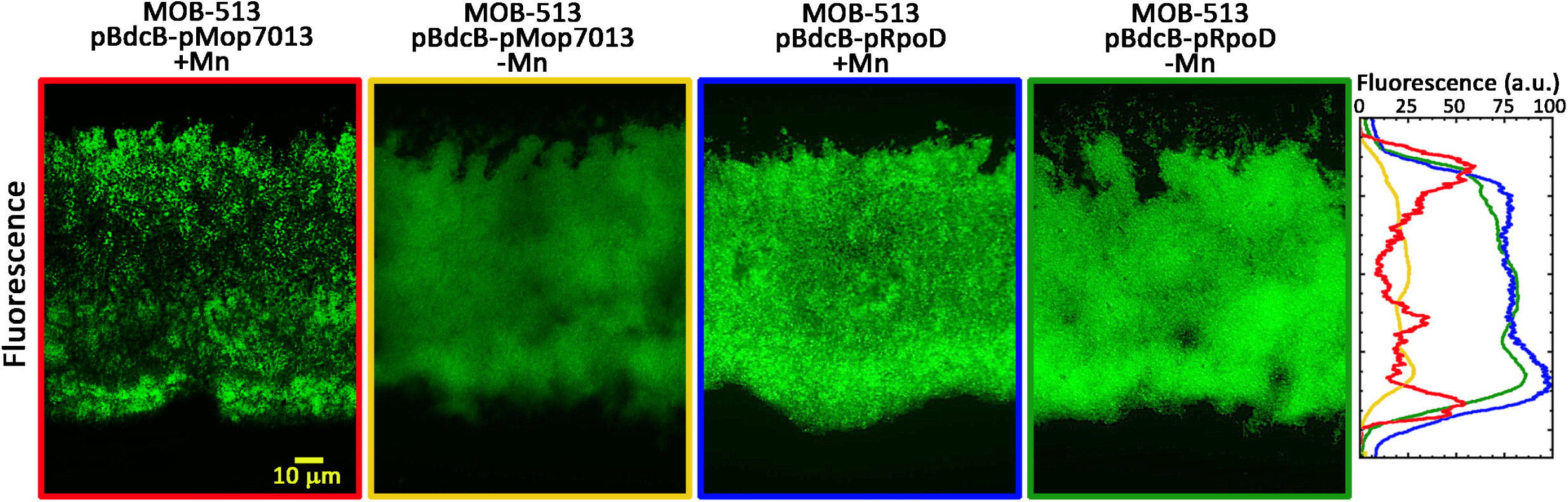
Analysis of the spatial expression of *mop7013* and *rpoD* genes in MOB-513-pBdcB macrocolony biofilms. Macrocolonies of MOB-513-pBdcB-pMop7013 and MOB-513-pBdcB-pRpoD were grown on Lept (-Mn) and Lept-Mn (+Mn) and thin-sectioned. Images are fluorescence micrographs that visualize the central region of cross-sections of 3-day-old macrocolonies. The spectral plot shows fluorescence of the *mop7013*::*gfp* and *rpoD*::*gfp* reporter fusions as a function of depth across the macrocolony cross-section. The colours of the spectra collate with the code of the reporter fusions and growth conditions indicated in the edges of the panels. Quantification of the spatial distribution of Gfp activities of reporter fusions across macrocolony cross-sections was performed using FIJI software. For each reporter fusion or growth condition, the highest fluorescence intensity value in the respective spectrum was arbitrarily (arbitrary units, a.u.) set to 100.

### High levels of c-di-GMP induce specific changes in the MOB-513 proteome

To more closely examine how the over-expression of *BdcB* accelerates and enhances Mn(II) oxidation and to identify proteins besides MopA involved in the MOB-513 Mn(II) oxidation process, label free quantification was conducted on proteomes from 2-day-old macrocolonies of MOB-513-pEmpty and MOB-513-pBdcB (Proteomic Supplementary Information).

Both Mop7013 and Mop7014 were detected only in strain MOB-513-pBdcB (Table S3), consistent with the higher expression levels of their genes in this strain compared to MOB-513-pEmpty. Interestingly, for MOB-513-pEmpty, a GGDEF/EAL-domain protein (Pres513_4541) showed increased levels in the presence of Mn(II) (Table S1). Furthermore, for MOB-513-pBdcB, a hypothetical PilZ-domain protein (Pres513_6471), was Mn(II) upregulated (Table S1). Together, these data confirm that both Mop7013 and Mop7014 are proteins upregulated by c-di-GMP and provide a molecular link between c-di-GMP signalling and Mn(II) oxidation.

### Overexpression of BdcB in MOB-513 improves lyophilisation and groundwater Mn(II) oxidation

Previous studies showed the usefulness of MOB lyophilised cultures to replace large volumes of inoculum and to enhance groundwater Mn removal performance (10). Since MOB-513-pBdcB shows a higher biofilm formation capacity and is a hyper-oxidant bacterial strain, we expanded our analysis to consider the biotechnological relevance of c-di-GMP function in MOB-513. We determined that high c-di-GMP levels improved the lyophilisation performance of MOB-513 (Table 2). In addition, for both MOB-513-pEmpty and MOB-513-pBdcB, the higher the initial content of BMnOx present in the cultures, the higher the observed survival rate (SR) (p<0.05) (Table 2). Next, the capacity of MOB-513-pEmpty, MOB-513-pBdcB and MOB-513-pBpdeA fresh cultures and lyophiles immobilized on sands to oxidize Mn(II) present in groundwater was analysed. Sand inoculated with MOB-513-pBdcB fresh cultures or lyophiles, both grown in Lept or Lept-Mn, showed a higher Mn(II) oxidation capacity than those inoculated with MOB-513-pEmpty (Table 2). On the other hand, and as previously observed (10), bacterial adaptation to Mn(II) performed by growing the strains in Lept-Mn enhanced the oxidation of Mn(II) present in groundwater (Table 2). No BMnOx was detected for MOB-513-pBpdeA at the assayed timepoints (Table 2).

**Table 2.** Quantification of BMnOx production, bacterial growth and lyophilisation survival ratios of MOB-513-pEmpty, MOB-513-pBdcB and MOB-513-pBpdeA static cultures grown in Lept and Lept-Mn and analysis of groundwater Mn(II) oxidation performed by bacteria inoculated sands. ^*^ND: Not detected.

| Quantification* | MOB-513-pEmpty |  | MOB-513-pBdcB |  | MOB-513-pBpdeA |  |
| --- | --- | --- | --- | --- | --- | --- |
|  | Lept | Lept-Mn | Lept | Lept-Mn | Lept | Lept-Mn |
| Initial BMnOx in the culture (µg/mL) | ND* | 0.44±0.08 | ND* | 10.45±0.30 | ND* | ND* |
| Fresh cultures (CFU/mL) | 2.50±0.27<br>x 10<sup>10</sup> | 2.25±0.24<br>x 10<sup>10</sup> | 6.50±0.24<br>x 10<sup>10</sup> | 2.00±0.22<br>x 10<sup>11</sup> | 1.65±0.22<br>x 10<sup>9</sup> | 2.55±0.22<br>x 10<sup>9</sup> |
| Lyophiles (CFU/mL) | 3.25±0.34<br>x 10<sup>07</sup> | 3.60±0.31<br>x 10<sup>07</sup> | 1.93±0.20<br>x 10<sup>08</sup> | 2.33±0.27<br>x 10<sup>09</sup> | 9.9±0.25<br>x 10<sup>5</sup> | 1.79±0.24<br>x 10<sup>6</sup> |
| SR (%) | 0.130 | 0.160 | 0.296 | 1.165 | 0.06 | 0.07 |
| BMnOx produced in groundwater by fresh cultures adhered to sands (µg/mL) | 1.25±0.10 | 6.72±0.25 | 15.98±0.33 | 23.92±0.41 | ND* | ND* |
| Groundwater BMnOx produced by lyophiles adhered to sands (µg/mL) | 1.28±0.09 | 6.46±0.31 | 17.90±0.35 | 24.10±0.42 | ND* | ND* |

## DISCUSSION

The presence of Mn(II) in groundwater impacts negatively on people unless it is appropriately treated. Biological sand filter technology is widely used for groundwater potabilization as an eco-friendly strategy that does not require chemical addition, increases groundwater treatment capacity, and reduces operative costs (2). Biofilters harbour microbial communities recruited by the groundwater to be treated or by bioaugmentation approaches (4–8). Bioaugmentation using bacteria with high biofilm-forming and Mn(II)-oxidising capabilities provides an inexpensive, simple and efficient strategy for immobilizing these bacteria in the sand filters and optimising Mn removal (4, 5, 8). Although many efforts have been made towards isolating bacteria with these characteristics (11), no studies have been performed to assess the importance of a biofilm lifestyle on the Mn(II) oxidation process.

The role of c-di-GMP in biofilm formation and in numerous bacterial functions such as motility, regulation of cell cycle, differentiation, and virulence (13) is well understood. In this work, we expand these to describe a novel role for this second messenger in the regulation of Mn(II) oxidation in the environmental *P. resinovorans* strain MOB-513, isolated from sand biofilters that currently remove groundwater Mn with high efficiency (11). Our data shows that high levels of c-di-GMP increase both biofilm formation and Mn(II) oxidation capacities in MOB-513. Moreover, c-di-GMP-driven oxidation occurs by enzymatic activity, and affects the abundance of Mop proteins previously linked to Mn(II) oxidation in other bacteria (20, 26, 30).

Biofilm growth promotes the formation of resource gradients and micro-niches that uniquely affect cellular physiology and that are not observed in liquid culture (29, 31). In macrocolony biofilms, oxygen becomes limited for cells at depth in the biofilm due to its consumption by those closer to the periphery, whereas the opposite happens with the nutrient gradient being more abundant when closer to the nutrient-providing agar. In addition to these previously observed gradients (29, 31, 32), we also saw evidence of stratified Mn(II) oxidation carried out by specific layers of cells in MOB-513 macrocolony biofilms. The spatial localization of BMnOx in the macrocolony biofilms has never been characterised before, and curiously, the levels of the second messenger c-di-GMP play a key role in defining when and where the microorganism carries out this oxidation within the biofilm. We found that Mn(II) ions are maximally oxidised at different depths: in the macrocolonies of MOB-513 control strain and the strain overexpressing *BpdeA*, BMnOx accumulation is only pronounced in the upper, more oxygenated portion of the biofilm, while in the strain overexpressing *Bdgc-B*, additionally the lower microoxic/anoxic portion also accumulates BMnOx.

Our results show that MOPs are the main enzymes responsible for Mn(II) oxidation in *P. resinovorans* MOB-513 biofilms and that the expression of these enzymes is positively regulated by c-di-GMP. The transcriptional profile of *mop7013* and *mop7014* genes also showed an increased expression in the presence of Mn. Accordingly, it has been observed an increment of the levels of the MopA protein in *A. manganoxydans* strain SI85-9A1 in the presence of the metal (26). Proteomic data showed that the abundances of Mop7013 and Mop7014 proteins were altered by this second messenger, and the areas in which Mn(II) oxidation takes place in MOB-513-pBdcB macrocolony biofilms spatially correlate with the expression of *mop7013* and *mop7014* when Mn(II) is present supporting their participation in the Mn(II) oxidation process.

High c-di-GMP levels and the presence of Mn(II) extensively remodelled MOB-513 proteome. Among the upregulated proteins for MOB-513 in the presence of Mn(II) we observed a GGDEF/EAL-domain protein, suggesting a role in changing the intracellular c-di-GMP levels in response to this metal. Furthermore, we identified a PilZ-domain protein that was upregulated for MOB-513-pBdcB in the presence of Mn(II). The PilZ domain is a c-di-GMP-binding protein domain (33). Single-domain PilZ proteins are widespread and several of them mediate cellular functions and bacterial behaviours known to be regulated by c-di-GMP (34). The upregulation of this protein in MOB-513-pBdcB in the presence of Mn(II) suggests that it may have a role in Mn(II) oxidation signalling by c-di-GMP. Future analysis of both proteins in MOB-513 biofilm formation and Mn(II) oxidation will clarify their role in the c-di-GMP signalling pathway mediating Mn(II) oxidation. Previous results in *P. putida* GB-1 demonstrated that deletion of the c-di-GMP binding master regulator *fleQ* results in increased levels of the enzyme MopA, while the activities of the other two MCOs, MnxG and McoA decreased (20). FleQ may be the regulator that links c-di-GMP signalling to Mn(II) oxidation processes, although further studies are necessary to understand this regulatory network.

Bacteria can use Mn(II) oxidation as an adaptive advantage to survive adverse environmental conditions (16). The second messenger c-di-GMP appears to be the link between biofilm-forming and Mn(II) oxidizing capacities of MOB-513, suggesting that both traits had co-evolved as an adaptation to the biofilm lifestyle. These two mechanisms may act simultaneously to some extent, with c-di-GMP determining the spatial and temporal distribution of BMnOx across the MOB-513 biofilm. Several advantages of possessing Mn(II) oxidizing activity for the biofilms may be considered. Consistent with previous reports indicating that Mn(II) oxidation increases tolerance to oxidative stress (35), Mn(II) oxidizing cells are able to resist the input of high concentrations of this metal present in the environment and after oxidation the ion becomes insoluble. Insoluble accumulations of BMnOx can prevent predation or viral attack (16), protect from radiation (36), and can enable the oxidative degradation of natural organic matter, with bacteria gaining energy from this process (37). Therefore, Mn(II) oxidation within biofilms may not only have a role as an antioxidant and provide a protective barrier against different stressors, but may also benefit the biofilm through the energy they acquire by detoxifying high concentrations of Mn(II) when present.

Finally, we expanded the analysis to consider the biotechnological relevance of c-di-GMP Mn(II)-oxidation control in MOB-513. We examined its impact on lyophilisation and the Mn(II)-oxidation efficiency of lyophiles, with our data showing that high levels of c-di-GMP correlate with higher lyophilisation efficiencies and higher groundwater Mn(II) oxidation capacities of MOB-513 lyophiles.

Several questions, currently under investigation, include: is c-di-GMP-regulated oxidation widespread among MOB? What are the cues controlling the expression of Mop proteins across biofilm depth in the presence of Mn in the culture medium? What are the main MOB-513 DGCs/PDEs controlling the c-di-GMP signalling? How do the differentially expressed proteins detected in our proteomic assays affect signalling and impact on Mn(II) oxidation? Overall, these results provide evidence to support the role of c-di-GMP in biofilm formation and Mn(II) oxidation in *P. resinovorans* and provide new strategies for optimising the biotechnological application of this bacterium in bioremediation.

## MATERIALS AND METHODS

### Plasmids, strains, and growth conditions

The plasmids and bacterial strains used in this study are listed in Table 3. Proteins BdcB (gene BB3903) and BpdeA (gene BB2664) from *Bordetella bronchiseptica* (22, 23), were overexpressed by using the vector pBBR1MCS-5 under the control of the constitute promoter *nptII* (38). These plasmids and the corresponding empty vector (pEmpty) were transformed into *P. resinovorans* MOB-513 by electroporation (39). The plasmid pWspR19 corresponds to the pBBR1MCS-5 vector containing a constitutively active allele of the DGC WspR (24). For the construction of promoter-*gfp* reporter fusions, the predicted promoters of genes *mop7013, mop7014*, and *rpoD* were cloned (primers described in Table 4) upstream the promoter-less *gfp* gene in the pPROBE-KT vector (40).

**Table 3:** Plasmids and strains used in this study.

| Strain or plasmid | Genotype | Antibiotic<sup>r</sup> | Reference |
| --- | --- | --- | --- |
| *Plasmids* |  |  |  |
| pEmpty | pBBR1MCS-5 (with *nptII* promoter) | Gm | (38) |
| pBdcB | pBBR1MCS-5 derived plasmid with *BdcB* inserted downstream the *nptII* promoter | Gm | (22) |
| pBpdeA | pBBR1MCS-5 derived plasmid with *BpdeA* inserted downstream the *nptII* promoter | Gm | (23) |
| pPROBE-KT | rep<sup>p</sup> BBR1 promoterless *gfp* | Km | (40) |
| p7013 | pPROBE-KT derived plasmid with the *mop7013* promoter inserted upstream the *gfp* gene | Km | This work |
| p7014 | pPROBE-KT derived plasmid with the *mop7014* promoter inserted upstream the *gfp* gene | Km | This work |
| pRpoD | pPROBE-NT derived plasmid with the *rpoD* promoter inserted upstream the *gfp* gene | Km | This work |
| *Strains* |  |  |  |
| *P. resinovorans* MOB-513 | Manganese oxidizer, wild type | Cm | (11) |
| MOB-513-pEmpty | MOB-513 containing pEmpty | Cm, Gm | This work |
| MOB-513-pBdcB | MOB-513 containing pBdcB | Cm, Gm | This work |
| MOB-513-pBpde | MOB-513 containing pBpdeA | Cm, Gm | This work |
| PAO1-pEmpty | *P. aeruginosa* PAO1 containing pEmpty | Gm | (24) |
| PAO1-pWspR | *P. aeruginosa* PAO1 containing pWspR19 | Gm | (24) |
| MOB-513-pBdgc-pPROBE-KT | MOB-513-pBdcB containing pPROBE-KT | Gm, Km | This work |
| MOB-513-pBdcB-pMop7013 | MOB-513-pBdcB containing p7013 | Gm, Km | This work |
| MOB-513-pBdcB-pMop7014 | MOB-513-pBdcB containing p7014 | Gm, Km | This work |
| MOB-513-pBdcB-pRpoD | MOB-513-pBdcB containing pRpoD | Gm, Km | This work |

**Table 4:**
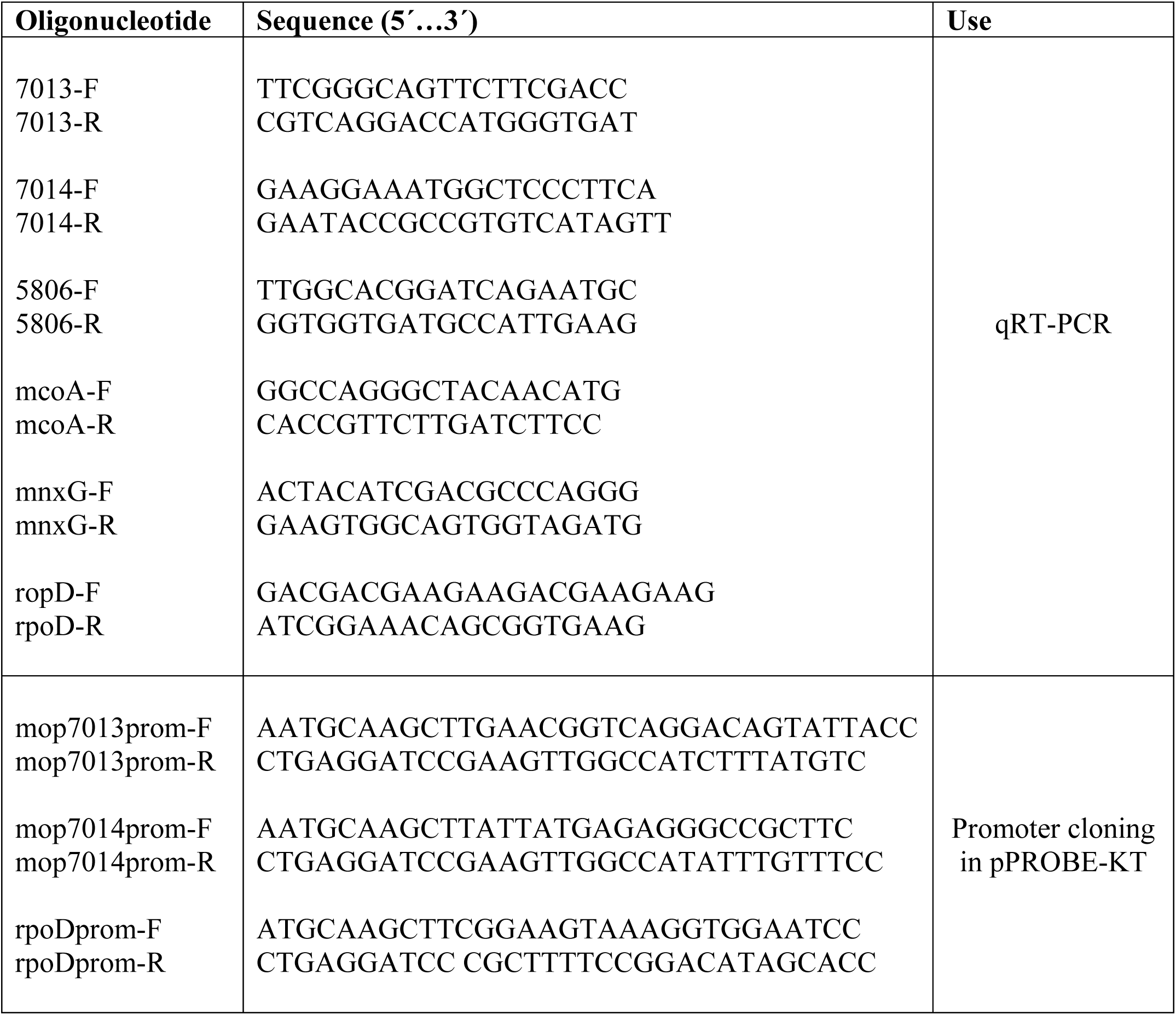
Oligonucleotides used in this study.

*P. resinovorans* MOB-513 strains were grown at 28°C in either Luria–Bertani (LB) or in Lept medium with or without supplementation of 100 µM MnCl_2_ (Lept-Mn or Lept) (41). *Escherichia coli* and *P. aeruginosa* strains were grown in LB medium at 37°C. When required, the following antibiotics at the specified concentrations were used: kanamycin (Km) 20 µg/mL, chloramphenicol (Cm) 10 µg/mL, and gentamicin (Gm) 10 µg/mL.

### Bacterial genome sequencing and bioinformatics techniques

*P. resinovorans* MOB-513 genomic DNA was sequenced using Illumina technology platform (Illumina Inc. USA) as previously described (42). The assembled genome of *P. resinovorans* MOB-513 was deposited at DDBJ/ENA/GenBank under the accession JAJOHH000000000.

### Quantification of intracellular c-di-GMP concentrations

The quantification of c-di-GMP levels was performed using the Cyclic-di-GMP Assay Kit from Lucerna (Catalog Number: 200-100). Cultures of the strains were adjusted to OD_600_ = 0.2 and set up for the assay in 30-µl aliquots along with assay reagents and serially diluted c-di-GMP standards, and then the c-di-GMP concentration was calculated according to the standard calibration curve. Appropriate sample dilution factors were multiplied to get the final c-di-GMP concentrations (pg/µL).

### Biofilm formation assays

Biofilm formation by MOB-513 strains was assessed as described previously (11). Macrocolony biofilms of MOB-513 strains were set up as previously described (25). Briefly, 5-µl of an overnight culture grown in LB and adjusted to an OD_600_ = 0.1 were spotted onto Lept or Lept-Mn agar plates and incubated at 28°C for up to 7 days.

### Motility assays

For each MOB-513 strain, 5-µl of an overnight culture adjusted to an OD_600_ = 1.5 was spotted onto the centre of LB swimming plates (0.3% agar) as previously described (43) and migration zone after 72 h of incubation were measured.

### BMnOx quantifications

Mn(II) oxidation was quantified by using Leucoberbelin Blue (LBB) dye solution as previously described (11).

### Cryosectioning of macrocolony biofilms and brightfield and fluorescence microscopy

The procedures for cryoembedding and cryosectioning macrocolony biofilms of MOB-513 strains into 5-µm-thin cross-sections were carried out as previously described (25). Five-micrometre-thick macrocolony sections perpendicular to the plane of the macrocolony were sectioned using a cryostat (Thermo Fisher Scientific) set at 20°C using disposable Sec35 blades (Thermo Fisher Scientific). The sections were placed on microscope slides and mounted with mowiol 4-88.

### Quantification of Mn(II) oxidase activity

Mn(II) oxidase activity of crude total protein extracts (PE) of MOB-513 strains was evaluated by quantifying the conversion reaction of Mn(II) to BMnOx *in vitro* using LBB (11). 1mL of initial reaction mixtures (10 mM HEPES pH 7.5, 0.020 mg/mL PE and 5 mM MnCl_2_) was incubated statically at 28°C for about 24 h. The effect of Ca(II) or Cu(II) was studied by adding 25 mM CaCl_2_ (26) or 1 mM CuSO_4_ (27), to the reaction mixtures. Control reaction mixtures were heated at 95°C for 15 min. Values of Mn(II) oxidase activity were normalized by the activity measured for each reaction mixture at t_0_, *i*.*e*., before the incubation at 28°C.

### RNA preparation and quantitative real-time PCR (qRT-PCR)

RNA was extracted from pools of macrocolonies using 1 ml TRIzol reagent (Invitrogen) according to the manufacturer’s instructions, subjected to DNAse (Promega) treatment and cDNA was synthesized using M-MLV RT (Promega, USA). Gene specific primers used (Table 4). Then, qRT-PCRs were performed in a Mastercycler realplex thermal cycler (Eppendorf) using Platinum Taq DNA polymerase (Invitrogen) and SYBR Green I (Roche) to monitor double-stranded DNA (dsDNA) synthesis. The *rpoD* gene was used as internal control (44).

### Quantification of Gfp fluorescence in macrocolony cells

Quantification of Gfp fluorescence in macrocolonies of MOB-513 co-expressing DGC-B, was assayed as previously described (45). Briefly, 2-days-old macrocolonies of MOB-513 strains grown on Lept or Lept-Mn were collected, washed, and resuspended in PBS buffer to achieve an OD_600_ = 0.5. Aliquots of 200 µL of each cell suspension were placed into 96-well flat bottom black plates (Greiner Bio-One). Fluorescence intensity (F) and the final OD_600_ of each sample was recorded on a Synergy 2 Multi-Mode Microplate Reader (BioTek) using excitation and emission filters with wavelengths of 485±20 nm and 535±20 nm, respectively. F values were normalized applying the following formula:

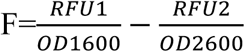

where RFU^1^ and OD^1^_600_ are the fluorescence intensity measured by instrument and the final OD_600nm_ determined for the strain expressing a promoter-*gfp* fusion, respectively. RFU^2^ and OD^2^_600_ are the same parameters determined for the sample of cells carrying the empty vector (pPROBE-KT).

### Proteomic assays - Label Free Quantification

Twelve macrocolony biofilms of MOB-513 strains were grown for 2 days on Lept or Lept-Mn agar plates. Then, they were scraped, resuspended in PBS, and subjected to protein extraction. Samples containing 30 µg of whole-cell proteins were sent to the Proteomics Core Facility of CEQUIBIEM where protein digestion and Mass Spectrometry analysis were performed by nano-HPLC coupled to a mass spectrometer with Orbitrap technology.

### Bacterial lyophilisation and quantification of bacterial survival ratios and BMnOx production

MOB-513 strains were lyophilized following as previously described (10). Viability of bacteria (CFU/mL) before and after lyophilisation was used to calculate the cell survival rates (SR) as follow:

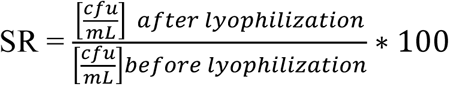

### Quantification of the oxidation of Mn(II) present in groundwater

The oxidation of Mn(II) present in groundwater carried out by sand-immobilized bacteria derived from fresh cultures and lyophiles was quantified as previously described (10).

### Statistical analysis

Quantifications were performed from three biological replicates and three technical replicates per sample. All data were statistically analysed using one-way ANOVA (p<0.05) or by Tukey’s test or by Student’s t-test (as indicated in Figure legends).

## ACKNOWLEDGMENTS

This work was supported by PICT2019-01081 and PIP-1516 from ANPCyT and CONICET, respectively. Work in the JGM lab was supported by BBSRC Grants BBS/E/J/000PR9797 and BB/T004363/1. AP’s short-research stay abroad was supported by IUBMB Wood-Whelan Fellowship. AP and LCC were fellows and FS, JF, JO, DOS and NG are staff members of CONICET. AP, DOS and NG conceived and designed the work, with contributions from FS, JF, JGM and JO. AP and LCC performed most of the experiments, with contributions from DOS and NG on macrocolony experiments. AP, DOS and NG wrote the paper, with contributions from FS, JF, JGM and JO.

## SUPPLEMENTAL MATERIAL

**FIG S1**. Subsystem classification of *P. resinovorans* strain MOB-513 genome properties by RAST annotation (https://rast.nmpdr.org/). The 7,489,538 bp genome of MOB-513 was sequenced and has a GC-content of 62.5 %, with 7,209 coding sequences. RAST annotation allowed the classification in subsystems. The most abundant categories are related to maintain basal cell functions such as: Amino Acids and Derivatives metabolism; Carbohydrates metabolism; Cofactors, Vitamins, Prosthetic Groups; Fatty Acids, Lipids, and Isoprenoids metabolism; Protein metabolism; Aromatic Compounds metabolism (that is a characteristic prominent metabolism in environmental *Pseudomonas*); DNA and RNA metabolism; Membrane Transport and Stress Response.

**FIG S2**. Analysis of the *mco* genes expression levels in the different strains. The macrocolonies of the strains MOB-513-pEmpty (A), MOB-513-pBdcB (B) and MOB-513-pBpdeA (C) were grown on Lept and Lept-Mn and subjected to qRT-PCR assays (as described in FIG 5) to analyse *mco3924* and mco*3296* gene expression. Bars indicate the expression levels of the genes in Lept-Mn (+Mn) relative to the expression levels at 2 days in the absence of Mn (-Mn). Values are the means of three biological replicates with three technical replicates each. Error bars indicate standard deviation. Data were analysed by Student’s t-test and asterisks indicate significant difference (p<0.05) between samples grown in the presence of Mn(II) and without this metal.

**FIG S3**. (A) Graphic of the MOB-513 chromosomal region centred on the focused genes *mop7013* and *mop7014* from *P. resinovorans* strain MOB-513. The online software Softberry (http://www.softberry.com/berry.phtml?topic=index&group=programs&subgroup=gfindb) was used for prediction of *mop7013* and *mop7014* promoter regions. The results predicted that these two genes are not part of an operon and each one has its own promoter. The figure shows each open reading frame as an arrow filled with different patterns and the size of the proteins as the number of amino acids. Arrows pointing to the right indicate that genes are on the forward strand and to the left that they are on the reverse one. Predicted function/description have been obtained from the RAST Server (https://rast.nmpdr.org/). (B) MOP7013 and MOP7014 from *Pseudomonas resinovorans* strain MOB-513 with predicted animal heme-peroxidase domains and Ca(II) binding regions labelled. Both, MOP7013 and MOP7014 contain two peroxidase domains related to the animal heme-peroxidase superfamily (PROSITE accession number PS50292), and following each of these domains are five and eleven hemolysin-type Ca(II) binding regions (PROSITE accession number PS00330), respectively.

**FIG S4**. Fluorescence quantification. Macrocolonies of the strains MOB-513-pBdcB-pRpoD, MOB-513-pBdcB-pMop7013 and MOB-513-pBdcB-pMop7014 were grown on Lept (-Mn) and Lept-Mn (+Mn) and Gfp fluorescence was quantified relative to Gfp of the MOB-513-pBdcB-pPROBE-KT strain. Quantifications were performed from three biological replicates and three technical replicates per sample and mean values (expressed in arbitrary units a.u.) and SD are presented in the figure.

**FIG S5**. Analysis of the spatial expression of *mop7013* and *rpoD* genes in MOB-513-pBdcB macrocolony biofilms. Macrocolonies of MOB-513-pBdcB-pMop7013 and MOB-513-pBdcB-pRpoD were grown on Lept (-Mn) and Lept-Mn (+Mn) and cryo-sectioned. Representative fluorescence and the corresponding merged phase-contrast/fluorescence (Merged) images of the mature central regions of the 3-day-old macrocolonies observed are shown. The spectral plot shows fluorescence of the *mop7013*::*gfp* and *rpoD*::*gfp* reporter fusions as a function of depth across the macrocolony, the highest fluorescence intensity value in the respective spectrum was arbitrarily (a.u.) set to 100.

**FIG S6**. Correlation between the oxidation of Mn(II) and c-di-GMP levels in macrocolonies of MOB-513. Left: representative images of 2-days old macrocolonies of MOB-513-pEmpty and MOB-513-pBdcB grown on Lept (-Mn) or Lept-Mn (+Mn) and the heatmap generated from proteomic data obtained for the respective macrocolonies. Rows and columns represent independent samples (3 biological replicates for each condition) and proteins, respectively. Red indicates higher protein abundance, while blue indicates lower protein abundance. The dendrogram represents results from hierarchical clustering and depicts similarities (Pearson correlation) between protein levels under the tested conditions. Cluster 1 contains 62 proteins that were downregulated with the overexpression of *dgc-B*, while cluster 2 contains 60 proteins that were upregulated with the overexpression of *dgc-B* (see Table S1 in the supplemental material).

## Proteomic Supplementary Information

Supplementary information of proteome analyses that includes Table S1-Table S5.

**Table S1**. KEGG pathway analysis of Down- and Up-regulated proteins in MOB-513-pEmpty and MOB-513-pBdcB in the presence of Mn(II).

**Table S2**. KEGG pathway analysis of cluster 1 and cluster 2 proteins in Figure S6.

**Table S3**. KEGG pathway analysis of differentially expressed ON proteins in MOB-513-pBdcB in the absence (-Mn) or the presence (+Mn) of manganese.

